# High efficiency organoid-derived cyst cultures as a drug discovery platform for polycystic kidney disease

**DOI:** 10.1101/2025.02.16.638545

**Authors:** Thitinee Vanichapol, Ruby Wangford, Misao Takemoto, Motonari Uesugi, Alan J Davidson, Veronika Sander

**Author notes:** Co-first authors (Thitinee Vanichapol, Ruby Wangford). Co-last authors (Alan J Davidson, Veronika Sander).

## Abstract

Autosomal dominant polycystic kidney disease (ADPKD) causes the slow, progressive formation of fluid-filled cysts within the kidney that ultimately leads to renal failure. Despite being the most common genetic kidney disease, the range of therapies for ADPKD is limited. Ureteric bud (UB) organoids derived from human pluripotent stem cells (hPSCs) have been used to model ADPKD. Here, we describe a protocol to generate UB organoids in large-scale suspension culture that is amenable to drug discovery. When differentiated from *PKD2*-deficient hPSCs, the UB organoids robustly form cysts within 7 days and reach nearly 100% cyst efficiency within ∼20 days. Cystic UB organoids express ADPKD markers and are responsive to cyst-reducing compounds, such as the BET inhibitor JQ1 and the NF-κB inhibitor QNZ, but not tolvaptan. As proof-of-principle of the platform as a drug discovery tool, we screened a small molecule library comprised of nutrient conjugates and identified M1, a palmitate-glucosamine-glycine conjugate, that reduced cyst formation. These results demonstrate that our scalable method of cystic UB organoid production provides a simple and highly efficient platform for ADPKD drug discovery.

**TRANSLATIONAL STATEMENT:** Autosomal dominant polycystic kidney disease (ADPKD) affects over 12 million people worldwide, yet the currently available therapy tolvaptan only moderately delays disease progression and is burdened with adverse effects. Improved treatments that are well tolerated and effective are urgently needed. We established a human ureteric bud organoid platform that enables ADPKD-like cysts to be cultured at large scale and low cost. These cystic organoids resemble human ADPKD cysts and are suitable for drug discovery. Identification of the nutrient conjugate M1 as a new lead molecule for reducing cyst formation highlights the clinical relevance of this model.

## INTRODUCTION

With a prevalence of 1 in 400 to 1 in 1000, autosomal dominant polycystic kidney disease (ADPKD) is the most common monogenic kidney disease and the most common genetic cause of kidney failure^1,2^. Mutations in the *PKD1* and *PKD2* genes account for 85% and 15% of ADPKD cases, respectively. *PKD1* and *PKD2* encode polycystin-1 (PC1) and polycystin-2 (PC2) transmembrane proteins that localize to the primary cilia of tubular epithelia. PC1 and PC2 form heterodimers that regulate calcium influx in response to tubular fluid flow. Mutations in either *PKD1* or *PKD2* lead to the disruption of the PC1/PC2 complex, causing aberrant calcium signaling alongside increased intracellular cAMP levels and multiple disease-mediating consequences that promote the formation of fluid-filled cysts that damage healthy tissue and impact kidney function. ADPKD manifests at an average age of 30-40 years, and over the following decades slowly progresses towards kidney failure^1,2^. ADPKD cysts develop from all segments along the nephrons and the collecting ducts. However, at later disease stages, cysts tend to accumulate at the collecting ducts where they become most damaging as they obstruct and compress upstream nephrons^3,4^. There is no cure for ADPKD, and the only FDA-approved treatment, the vasopressin V2 receptor inhibitor tolvaptan, only moderately slows disease progression. Patients must take tolvaptan twice daily for decades of life, which requires close monitoring and increases susceptibility to liver damage^5^. The dependence on this suboptimal yet high-cost therapy demands the development of more effective and better tolerated treatment options for ADPKD.

In recent years, hPSC-derived kidney organoids have been established for modeling polycystic kidney disease (PKD). These organoids are made from either gene-edited or patient-derived hPSCs and carry mutations in common cystogenic genes, including *PKD1* and *PKD2* for ADPKD, and *PKHD1* for autosomal recessive PKD (ARPKD). Depending on the mutation and on the protocol used to generate the organoids, cysts form in the organoids with variable efficiencies and either develop spontaneously^6–8^ or require stimulation with cAMP or forskolin^9–11^. Most studies so far have used the conventional nephron-containing kidney organoids (‘nephron organoids’), where cysts typically originate from the proximal and/or distal segments of the nephrons^6,8–13^.

Several promising drug screening approaches have been performed using human PKD-mimicking cystic organoid models. Tran *et al*.^6^ screened a kinase inhibitor library for cyst-reducing compounds and identified quinazoline (QNZ), a NF-κB inhibitor that blocks cyst formation in *PKD1-* and *PKD2*-deficient nephron organoids. Using a range of ADPKD- and ARPKD-mutation carrying organoids, Liu *et al*.^10^ found that cilium-autophagy signaling is linked to cystogenesis, and that treatment of the organoids with the autophagy inducer minoxidil attenuated cyst formation *in vitro* and in organoid xenografts in mice. Hiratsuka *et al*.^13^ used advanced flow culture to show more clinically relevant cyst formation from distal nephrons in *PKHD1*-mutant organoids. Furthermore, they discovered that flow-induced mechanical stress activates RAC1 and FOS, with cyst reduction achieved by targeting these signals with r-naxprofen and T-5224. Most recently, Vichy *et al*.^14^ used CRISPR base editing to recapitulate common ADPKD-causing mutations in nephron organoids and demonstrated the therapeutic potential of aminoglycoside drugs.

Nephron organoids lack ureteric bud (UB)-derived tissues and collecting ducts and are thus of limited use for modeling the sites of most severe cyst formation in ADPKD. In contrast, UB organoids are comprised of the renal ureteric epithelia that further differentiate into the branching collecting ducts^15–18^. UB organoids generated from hPSCs that carry cystogenic mutations therefore form cysts of ureteric origin^7,19^. A recent study by Mae *et al*.^20^ demonstrated the potential of cysts dissected from collecting duct organoids for drug discovery and identified retinoic acid receptor agonists as potential therapeutics to suppress cystogenesis.

Our lab has previously developed a protocol for human iPSC-derived nephron organoid production in large scale^21^. Here, we describe a method for generating UB organoids from nephron organoids, taking advantage of the plasticity of the distal tubule cells to be transdifferentiated into ureteric epithelia^7^. Both types of organoids are grown in suspension culture, allowing for scalability of the assays. We show that *PKD2*-deficient UB organoids develop into large cysts with high efficiency and are useful for drug testing and small molecule library screening.

## METHODS

### Induced pluripotent stem cell (iPSC) culture and manipulation

Human iPSC lines were maintained in mTeSR1 on geltrex-coated tissue culture plates, as described^22^. The MANZ-2-2 iPSC line^23^ was used for generating wildtype organoids. A *PKD2* loss-of-function mutation was introduced into MANZ-2-2 cells using CRISPR/Cas9 and a guide RNA targeting exon 1 of the *PKD2* gene (5’-GCGTGGAGCCGCGATAACCCCGG-3’^8^). Reverse transfection, cell sorting and screening for edited clones were performed as described^21^.

### Organoid generation

Nephron organoids were generated in suspension cultures as previously reported^22^. The protocol for UB organoid differentiation was modified from Howden *et al*^7^. In detail, ∼50 day 12 nephron organoids were washed once in DPBS, then digested in 1:1 TrypLE Express (Gibco)/Accutase (STEMCELL Technologies) for 15 min at 37°C. The organoids were gently triturated using a P1000 pipette until no clumps were visible. The cell suspension was passed through a 40 µm cell strainer to achieve a single cell suspension, diluted with 3 ml of Essential 6^TM^ medium (Gibco), then centrifuged 300g for 5 min. The cells were counted and resuspended in UE medium (Essential 6 medium supplemented with 100 ng/ml GDNF, 200 ng/ml FGF2 (both Peprotech), 2 µM CHIR99021 (STEMCELL Technologies) and 2 µM All Trans Retinoic Acid (ATRA, Sigma)) plus 10 µM Y-27632 (STEMCELL Technologies), then distributed into 2 wells of an ultra-low attachment 6-well plate (Corning), in 2 ml medium per well, at a density of 4-5×10^5^ cells per well. The plates were left undisturbed for 72 hrs. On day 3, a half medium change was performed by aspirating 1 ml per well and adding 1 ml freshly prepared UE medium (without Y-27632). The plates were placed on a rocker (15° tilt, 25 rpm) from day 3 on. Another half medium change was performed on day 6. UB organoids were used for drug screening from day 9, for which the compounds were added to Essential 6 base medium (without UE factors and Y-27632).

### Compound screening

Methylcellulose embedding was adapted from Tran *et al*^6^. Briefly, 1.5 g autoclaved methylcellulose powder (Sigma) was dissolved in 45 ml of Essential 6 medium at 60°C. 5 ml of Essential 6 medium + 1x Glutamax (Gibco) and 1% Penicillin-Streptomycin (Gibco) were added to achieve a stock concentration of 30 µg/ml. The methylcellulose stock was centrifuged at 4000g for 2 hrs. For UB organoid embedding, the methylcellulose stock was diluted 1 in 4 in Essential 6 medium + 1x Penicillin/Streptomycin, + 25 µg/ml Plasmocin (Invivogen). 80 µl of diluted methylcellulose were added to each well of a flat bottom 96-well plate. 10 µl of day 9 *PKD2*^*-/-*^ UB organoids (in Essential 6 medium) were added to each well. The plate was then placed in an 37°C incubator for 1 hr for the methylcellulose to solidify. The compounds of the nutrient conjugate library were added at 10 µM in 10 µl Essential 6 medium, to achieve a final concentration of 1 µM. Screening was performed in triplicate wells, with ≥20 cystic organoids per well and DMSO as a negative control. Brightfield images of each well were taken on days 0, 3, 7 and 10 of each assay, and fresh compounds were added on days 3 and 7. Hit compound M1 was validated in a second round of 96-well triplicate assays, as well as in suspension UB culture in an ultra-low attachment 24-well plate, in 1 ml Essential 6 medium and ≥100 cystic organoids per well. (+)-JQ-1, QNZ (EVP4593), tolvaptan (all Sigma), T-5224 (Selleckchem) and TAK-242 (Cayman Chemical) were also screened in 24-well suspension assays.

### Toxicity assays and immunostainings

For assessing toxicity, compounds were added to Stage II medium of day 12 nephron organoids. Visual analysis for necrosis, shedding of dead cells, and debris in the media was performed after 3 days, as described^24^. Organoids were fixed in 4% paraformaldehyde for 20 min at RT. Paraffin sections and immunohistochemistry were performed as described previously^21,22^. The Click-iT™ Plus TUNEL Assay (Invitrogen) was used for assessing cell death on nephron organoid sections. Imaging was performed on a Zeiss LSM710 confocal microscope. For quantification of cell death, sections through the middle of ≥10 individual organoids were measured for total TUNEL+ cell area and normalized to Hoechst+ area using Fiji. Antibodies used were: PAX2 (BioLegend 901001), CDH1 (BD Biosciences 610181), RET (Novus NBP184568), GATA3 (Thermo MA1-028), KRT8 (DSHB TROMA), EPCAM (Sigma HPA026761).

### qPCR

Total RNA extraction was performed using the GENEzol TriRNA Pure Kit (Geneaid). RNAs from human fetal (spontaneously aborted, 30 weeks) and adult (a mix of 4 donors aged 62-83) kidney tissue were purchased from Takara. cDNA synthesis was performed using the High-Capacity RNA-to-cDNA™ Kit (Applied Biosystems). qPCRs were performed using the PowerUp^TM^ SYBR^TM^ Green PCR Master Mix (Applied Biosystems) and run on a QuantStudio 6 Flex RealTime PCR machine (Applied Biosystems). Primer sequences are listed in Table S1.

### Cyst measurements

The formation and growth of cysts was tracked using an EVOS inverted microscope. The percentage of cystic organoids was measured by counting the number of cystic organoids and free-floating cysts compared to the overall count of organoids and free-floating cysts. The cyst diameters were measured using Fiji.

### Statistical analysis

Significance was determined using one-way or two-way ANOVA or unpaired *t-*test in Prism (GraphPad). *P* values of ≤0.05 were considered statistically significant.

## RESULTS

### Large-scale production of UB organoids from nephron organoids

We have previously developed a protocol for differentiating kidney organoids from human iPSCs that uses suspension culture, enabling rapid and large-scale organoid production^21,22^. These kidney organoids contain segmented nephrons, podocyte clusters, stromal and endothelial cells, but lack ureteric epithelia-derived tissues, i.e. collecting ducts. We therefore refer to our original kidney organoids as ‘nephron organoids’. To test if our nephron organoids could induce a ureteric cell fate, we adapted the method of transdifferentiation of ureteric epithelium from the distal nephron segments by Howden *et al*.^7^ to suspension culture (Figure 1A). Day 12 nephron organoids were digested into single cells and replated in ultra-low attachment 6-well plates in serum-free Ureteric Epithelium (UE) medium^25^, containing Wnt-analogue CHIR99021, FGF2, retinoic acid and the key factor for UB induction, GDNF (Figure 1B, C). The cultures were left undisturbed for 72 hrs, resulting in the formation of small cell aggregates. From day 3 on, the cultures were placed on a rocking shaker to prevent clumping of the aggregates. Over the course of a week, the aggregates grow into UB organoids, elongate and exhibit internal tubular structures (Figure 1D). Starting from ∼50 nephron organoids that correspond to 8.9×10^5^±2.0×10^5^ single cells after dissociation, we can generate a yield of up to 2000 UB organoids (average 1292±561, counted on day 7; Figure S1). We next assessed marker gene expression in UB organoids versus nephron organoids. We find that the podocyte marker nephrin (*NPHS1*) and the proximal tubule marker cubilin (*CUBN*) are selectively expressed in nephron organoids, while distal nephron markers *GATA3* and *PAX2* are strongly enriched in the UB organoids. This result is consistent with a loss of nephron segments and a shift to a distal and ureteric fate in UB organoids. Furthermore, specific ureteric epithelia markers, including *WNT9B, WNT11, RET, KCNN4, KANK4, MMP7, SLC52A3* and *SLC9A2*^7^, are significantly enriched in the UB organoids (Figure 1E). Immunolabeling of paraffin sections of UB organoids revealed protein expression of CDH1, PAX2, RET, GATA3 and KRT8, further highlighting their distal nephron / UE identity (Figure 1F). Taken together, we demonstrate that the transdifferentiation of nephron organoids into UB organoids can be achieved at scale in suspension.

**Figure 1.**
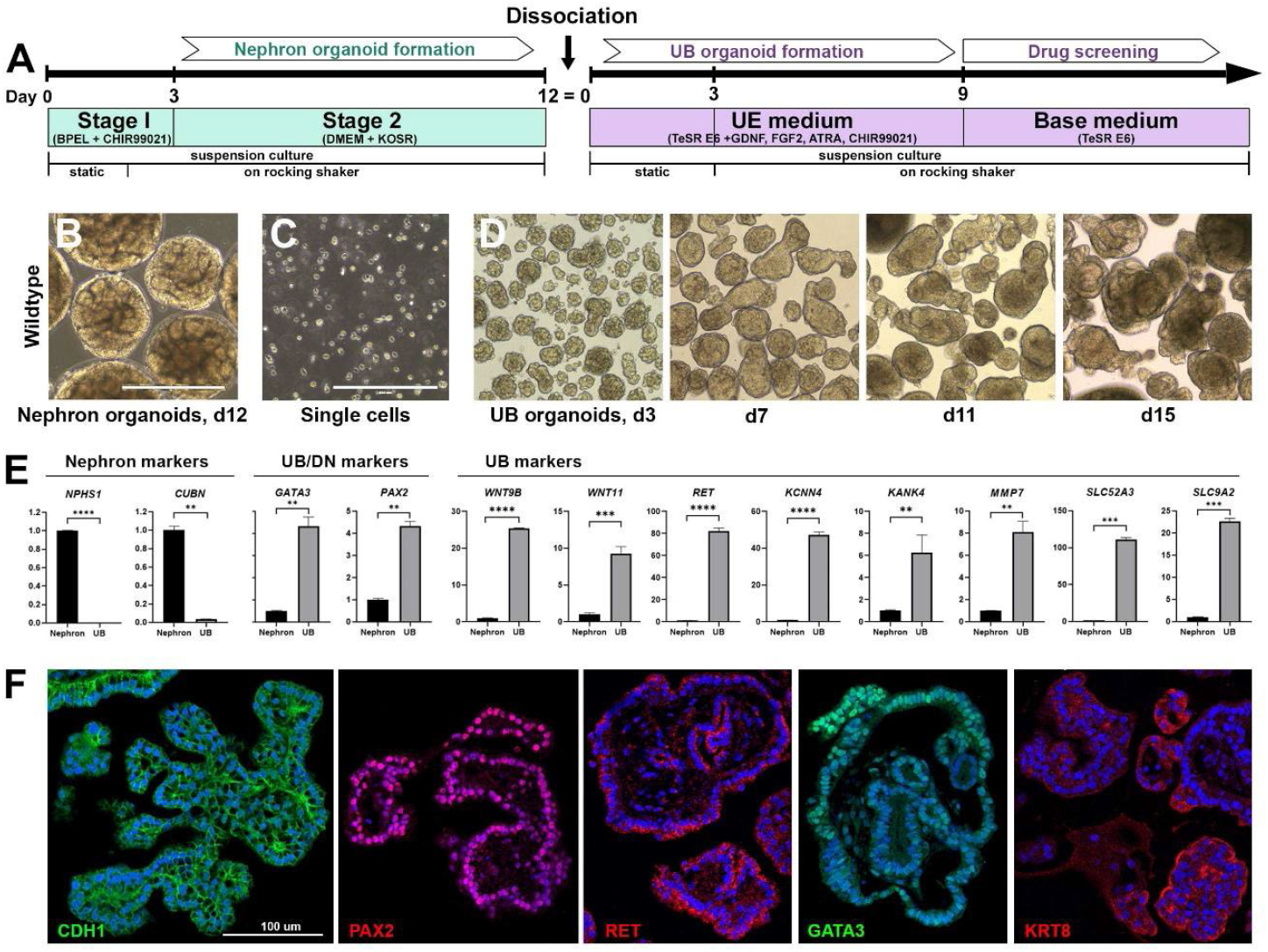
Ureteric bud organoid generation from nephron organoids. **A**. Schematic of the protocol. **B**. Day (d) 12 nephron organoids differentiated from wildtype iPSCs. **C**. Enzymatic digestion of whole nephron organoids into a single cell suspension, marking d0 of UB differentiation. **D**. Development of UB organoids over 15 days. **E**. qPCR analysis comparing d12 nephron organoids and d7 UB organoids for nephron and UB marker gene expression. ** p≤ 0.01; *** p≤ 0.001; **** p≤ 0.0001 (Unpaired *t*-test). **F**. Immunohistological characterization of UB organoids showing expression of distal tubule (CDH1, PAX2, GATA3) and UB markers (RET, KRT8). Nuclear counterstain, Hoechst. Scale bars: 400 µm (B-D); 100 µm (F).

### *PKD2*-deficient UB organoids develop into rapidly expanding cyst cultures

To apply our suspension UB organoid protocol to model ADPKD cysts, we used CRISPR/Cas9 gene editing to disrupt the *PKD2* gene, previously shown to lead to cyst formation in nephron organoids^8,26^. Nephron organoids derived from biallelic *PKD2* mutant (*PKD2*^*-/-*^) iPSC clones develop indistinguishably from wildtype nephron organoids (Figure S2A) before spontaneously forming cysts at around day 20. The cysts increase in number and size yet remain at a relatively low frequency of 13.4±5.7%, compared to 0.9±0.6% cysts formed in isogenic control nephron organoids (measured on day 26, Figure S2B). UB organoids differentiated from *PKD2*^*-/-*^ nephron organoids also resemble wildtype UB organoids in terms of size, shape and tubular composition over the first week of development, yet in contrast to nephron organoids, *PKD2*^*-/-*^ UB organoids spontaneously form cysts as early as day 5 (corresponding to day 17 overall; Figure 2A, B). Between day 7 and day 15 of UB organoid differentiation, cyst formation increases rapidly in these organoids and reaches 95% by day 25 (Figure 2A, B). The cysts are predominantly spherical, and over the course of 25 days their average diameter reaches 222 µm, with the largest cysts measuring >1 mm in diameter (Figure 2C). Multiple cysts appear to bud from a single UB organoid and often detach, resulting in free-floating cysts. In a single well of a 6-well plate, the number of cystic structures reaches an average of ∼460 per well, compared to ∼270 mostly non-cystic UB organoids in a wildtype assay (counted on day 25; Figure 2D, Figure S2C).

**Figure 2.**
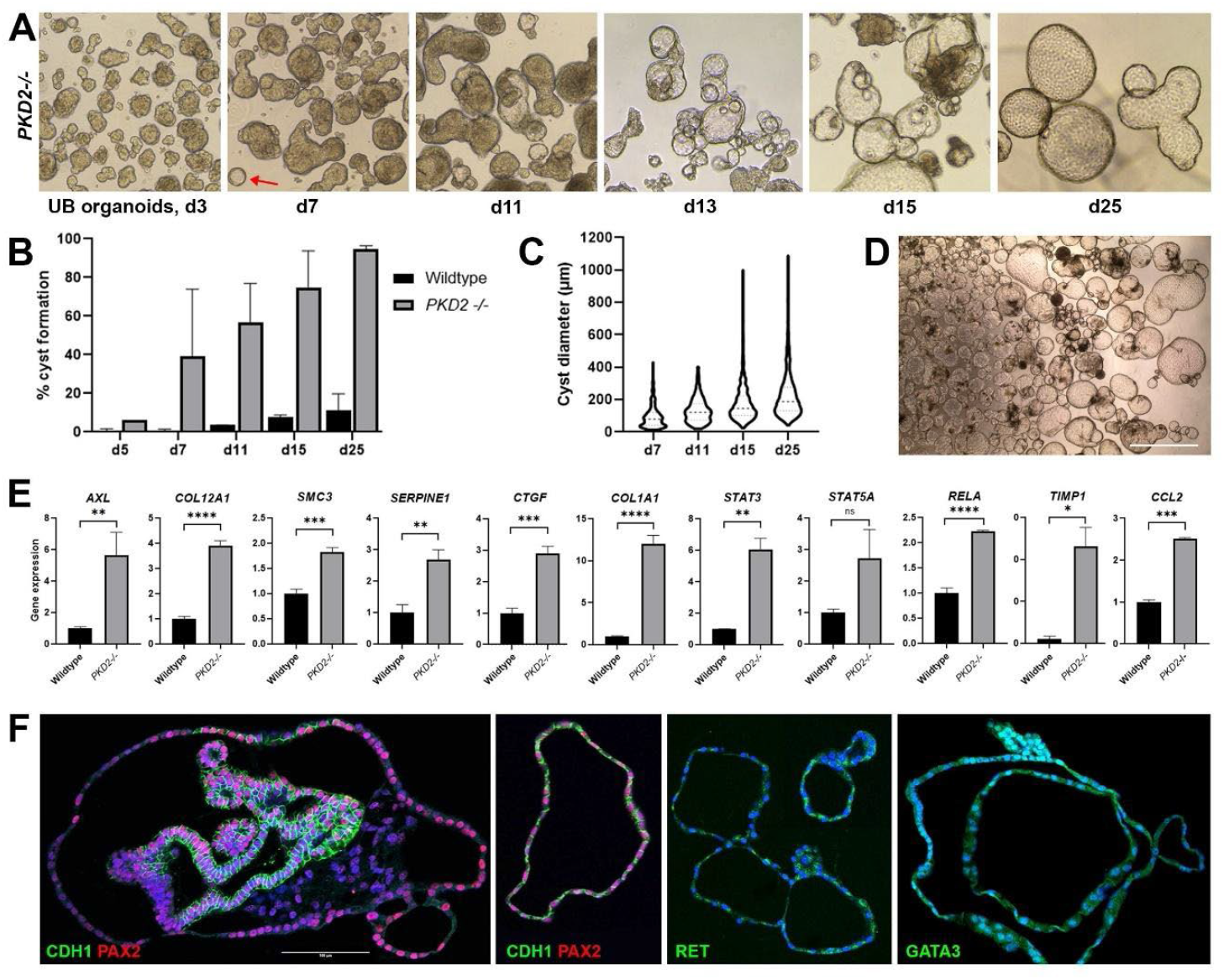
*PKD2*^*-/-*^ UB organoids form large-scale cyst cultures. **A**. Development of *PKD2*^*-/-*^ UB organoids over 25 days. Arrow shows an early cyst on d7. **B**. Quantification of cyst development in wildtype and *PKD2*^*-/-*^ UB organoids. **C**. *PKD2*^*-/-*^ cyst diameters increase over time. **D**. Example of a high efficiency cyst culture at d17. **E**. Upregulation of ADPKD marker genes in *PKD2*^*-/-*^ compared to wildtype UB organoids. ** p≤ 0.01; *** p≤ 0.001; **** p≤ 0.0001 (Unpaired *t*-test). **F**. Immunohistological analysis of *PKD2*^*-/-*^ UB organoids, showing expression of distal tubule and UB markers CDH1, PAX2, RET and GATA3 in the cystic epithelium. Nuclear counterstain, Hoechst. Scale bars: 400 µm (A); 1000 µm (D); 100 µm (F).

We next characterized the cyst cultures for expression of genes involved in ADPKD pathogenesis^27,28^, including effector genes of YAP1 signaling (*CTGF, SERPINE1, SMC3, AXL*), JAK-STAT signaling (*STAT3, STAT5A*), fibrosis-associated genes (*COL1A1, COL12A1, TIMP1, TGFB2*), monocyte attractant chemokine *MCP1*/*CCL2* and inflammation-associated NF-κB component *RELA*. All these marker genes were upregulated between 0.8-to 22-fold in the cystic *PKD2*^*-/-*^ UB organoids compared to wildtype UB organoids (Figure 2E). Immunohistochemistry revealed that the cysts comprise a single epithelial layer that labels for the distal nephron / UB markers CDH1, PAX2, RET and GATA3 (Figure 2F). Proximal tubule markers LTL and CUBN and the podocyte marker PODXL are not detectable in the cystic UB organoids (not shown). Based on these findings, we conclude that we have developed a new scalable method for producing cystic organoids with high efficiency.

### A platform for drug screening

Taking advantage of the scalability of our cyst cultures, we explored whether the platform has utility for drug discovery. We designed an assay to monitor cyst growth over 10 days, starting on day 9 of UB organoid development when the cysts are readily detectable, yet still relatively small (∼100 µm; Figure 2C). We used 24-well ultra-low attachment plates with ≥100 organoids in 1 ml of base medium per well with compounds added on day 0 of the assay and replaced with fresh compound with the medium changes on days 3 and 7 (Figure 3A). All organoids per well were captured by brightfield imaging on days 0, 3, 7 and 10, allowing quantitation of cyst number and diameter.

**Figure 3.**
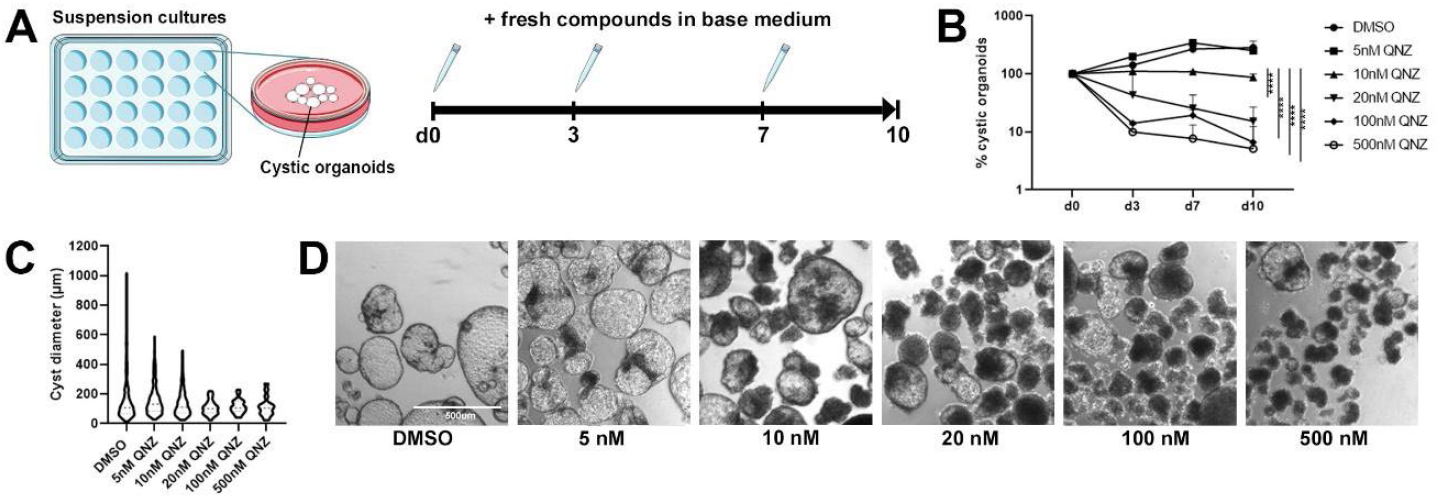
QNZ treatment reduces cyst numbers and sizes. **A**. Schematic and timeline of the assay. **B**. Numbers of cystic organoids (expressed as % of all organoids) after treatment with DMSO and different concentrations of QNZ over 10 days. **** p≤ 0.0001 (two-way ANOVA with Tukey’s multiple comparisons test). **C**. Cyst diameters after treatment with DMSO and QNZ on day 10. **D**. Representative brightfield images of DMSO- and QNZ-treated UB *PKD2*^*-/-*^ organoids on day 10. Scale bar, 500 µm.

We used the NF-κB inhibitor QNZ as positive control, based on recent work by Tran *et al*.^6^ where ≥5 nM QNZ was found to potently inhibit cyst growth and expansion in nephron organoids. We tested a range of QNZ concentrations from 5 nM to 500 nM and quantitated the percentage of cystic organoids among the total number of UB organoids. In our assay, 5 nM QNZ did not inhibit cyst formation, however there was a significant reduction in cyst number, as early as day 3, with concentrations ≥10 nM (Figure 3B). A direct comparison of the cyst diameters on day 10 did not reveal a significant difference in average cyst size between controls and QNZ-treated cystic organoids. However, a concentration-dependent decrease in very large cysts, i.e. cysts ≥200 µm was observed (Figure 3C). QNZ-treated cystic organoids appeared necrotic by brightfield microscopy (Figure 3D). Similarly, treatment of wildtype nephron organoids with QNZ ≥10 nM led to severe toxicity and necrosis, evidenced by a ∼4.5-fold increase of TUNEL-labeled dying cells at concentrations of 10 nM or 20 nM of QNZ, compared to DMSO-treated controls (Figure S3A-C). Taken together, these observations confirmed the cyst-reducing effect of QNZ and suggest that QNZ acts to inhibit cyst initiation and large cyst expansion by inducing cell death. The effect of QNZ on nephron organoids is of concern, particularly as we found that this occurs at a ∼1000-fold lower concentration than reported^6^, and suggests that it causes general toxicity to healthy kidney tissue. We also tested the c-FOS/activator protein (AP)-1 inhibitor T-5224^29^ and found that in our assay it had no effect on cyst numbers and instead it increased cyst size at ≥5 µM (Figure S4), contrary to the size-reducing effect shown in distal nephron cysts of *PKHD1*^-/-^ organoids under flow^13^. T-5224 was not toxic to nephron organoids at 20 µM (Figure S3).

### Tolvaptan does not affect cyst growth

We next tested the FDA-approved ADPKD drug tolvaptan, an inhibitor of vasopressin receptor V2 (encoded by the *AVPR2* gene). Previous nephron organoid-based ADPKD models have failed to achieve cyst reduction upon tolvaptan treatment, presumably due to the lack or low level of *AVPR2* expression. We performed qPCR analysis to evaluate *AVPR2* levels in our nephron and UB organoids, both wildtype and *PKD2*^*-/-*^, as well as human fetal and adult kidney samples. This revealed that *PKD2*^*-/-*^ UB organoids had the highest *AVPR2* expression amongst the organoids, yet this was still ∼3 times lower than in fetal and adult kidneys (Figure 4A). Nevertheless, we treated the cystic *PKD2*^*-/-*^ UB organoids with a concentration range of tolvaptan (20 nM to 10 µM) and found that neither the number of cysts nor the average cyst sizes were significantly reduced at any of the concentrations tested (Figure 4B-D). Tolvaptan was also not toxic to nephron organoids (Figure S3).

**Figure 4.**
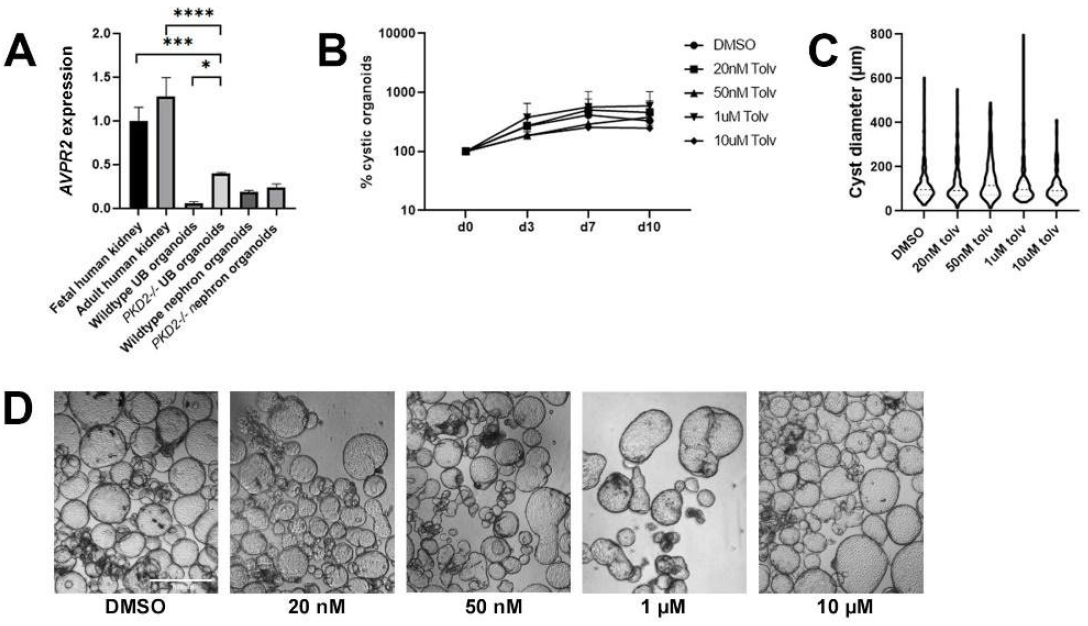
Tolvaptan does not affect cyst growth. **A**. Expression of vasopressin V2 receptor *AVPR2* in different organoids models and human kidneys. * p≤ 0.1; *** p≤ 0.001; **** p≤ 0.0001 (one-way ANOVA with Tukey’s multiple comparisons test). **B**. Percent of cystic organoids after treatment with DMSO and different concentrations of tolvaptan over 10 days. **C**. Cyst diameters after treatment with DMSO and tolvaptan on day 10. **D**. Representative brightfield images of DMSO- and tolvaptan-treated UB *PKD2*^*-/-*^ organoids on day 10. Scale bar, 500 µm.

### BET inhibitor JQ1 inhibits cyst formation

Inhibition of the BET bromodomain (BRD) protein Brd4 with the first-generation pan-BET inhibitor JQ1 has previously been shown to reduce cyst formation in two PKD mouse models^30^. More recently, a drug screen using cystic mini organoids derived from mouse *Pkd2*-mutant nephron progenitor cells identified four BET inhibitors (Bromosporine, BI-2536, AZD 5153, and CPI-203) that reduced organoid cyst formation^12^. We tested JQ1 in our platform at 0.1, 1 and 5 µM and found inhibition of cyst formation at all doses, with a significant reduction in cyst numbers observed at 5 µM (Figure 5A). Similar to QNZ, while the average cyst size did not change with JQ1 treatment, there was a dose-dependent reduction in large cysts (Figure 5B, C). No toxicity was observed in nephron organoids treated with JQ1 (Figure S3), suggesting that JQ1 specifically targets cystic UB epithelia.

**Figure 5.**
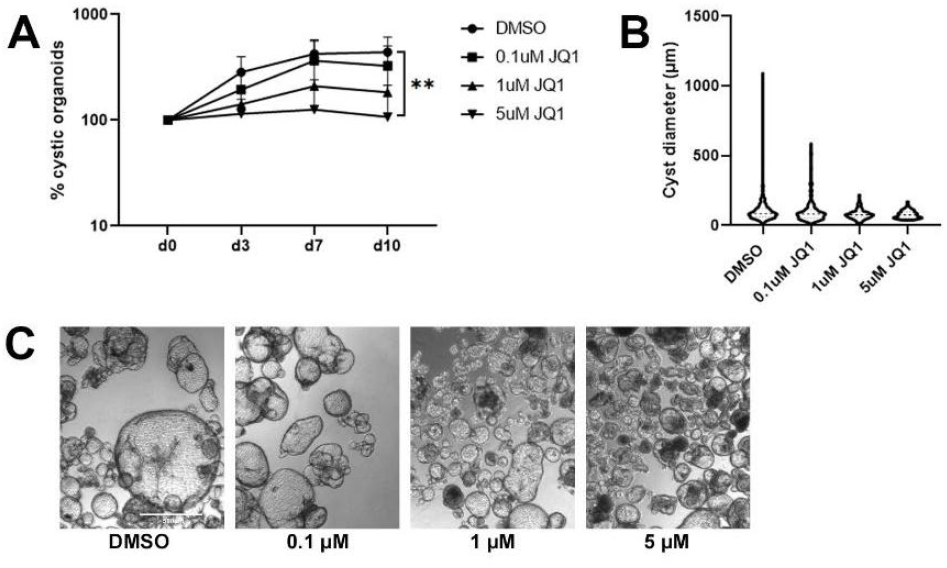
BET inhibitor JQ1 inhibits cyst formation and expansion. **A**. Percent of cystic organoids after treatment with DMSO and different concentrations of JQ1 over 10 days. ** p≤ 0.01 (two-way ANOVA with Tukey’s multiple comparisons test). **B**. Cyst diameters after treatment with DMSO and JQ1 on day 10. **C**. Representative brightfield images of DMSO- and JQ1-treated UB *PKD2*^*-/-*^ organoids on day 10. Scale bar, 500 µm.

### Screening of a novel drug library identifies compound M1

We next tested if our cystic UB organoid platform is suitable for larger scale drug screening. To do this, we screened a library composed of 308 covalently conjugated small molecule nutrients, including fatty acids, glucosamine, vitamins, amino acids, and biologically active peptides^31^. The concept of this library is that nutrient conjugates may combine or modulate the physiological functions of the individual nutrients, thus target metabolic or signaling pathways more potently and/or more specifically than the individual nutrients alone^32,33^. To accommodate testing the library in a low volume and high throughput format, we modified the assay to use 96-well plates and embedded the cystic *PKD2*^*-/-*^ UB organoids in methylcellulose, as described^6^. The compounds were initially evaluated in triplicate at 1 µM concentration in a volume of 10 µl, with ≥20 UB organoids per well, for the same timeline of 10 days as used for the suspension assays (Figure 6A). From this library we identified a single compound, M1, a conjugate of palmitate, glucosamine and glycines (Figure 6B), that is able to abolish cyst formation at 1 µM and significantly inhibit cyst size and cyst number at 0.5 µM (Figure 6C, D). At a concentration of 1 µM, M1 was clearly toxic to UB organoids (Figure 6E), although it did not affect nephron organoids suggesting it may selectively target cyst cells (Figure S3). The chemical structure of M1 closely resembles N-palmitoyl-D-glucosamine, a natural bacterial molecule shown to antagonize Toll-like receptor 4 (TLR4) and exert anti-inflammatory properties in mice^34,35^. We measured *TLR4* expression in the cystic *PKD2*^*-/-*^ UB organoids and found a significant 2-fold upregulation compared to wildtype UB organoids (Figure 6F), suggesting a role of TLR4 signaling in organoid cystogenesis. To test this hypothesis, we used the TLR4 inhibitor TAK-242^36^, previously described to ameliorate sepsis-induced acute kidney injury in sheep^37^, but found to be ineffective in a human clinical trial for the treatment of severe sepsis^38^. We found that TAK-242 dose-dependently decreased cyst numbers and cyst size (Figure 6G-I) and was non-toxic on nephron organoids (Figure S3).

**Figure 6.**
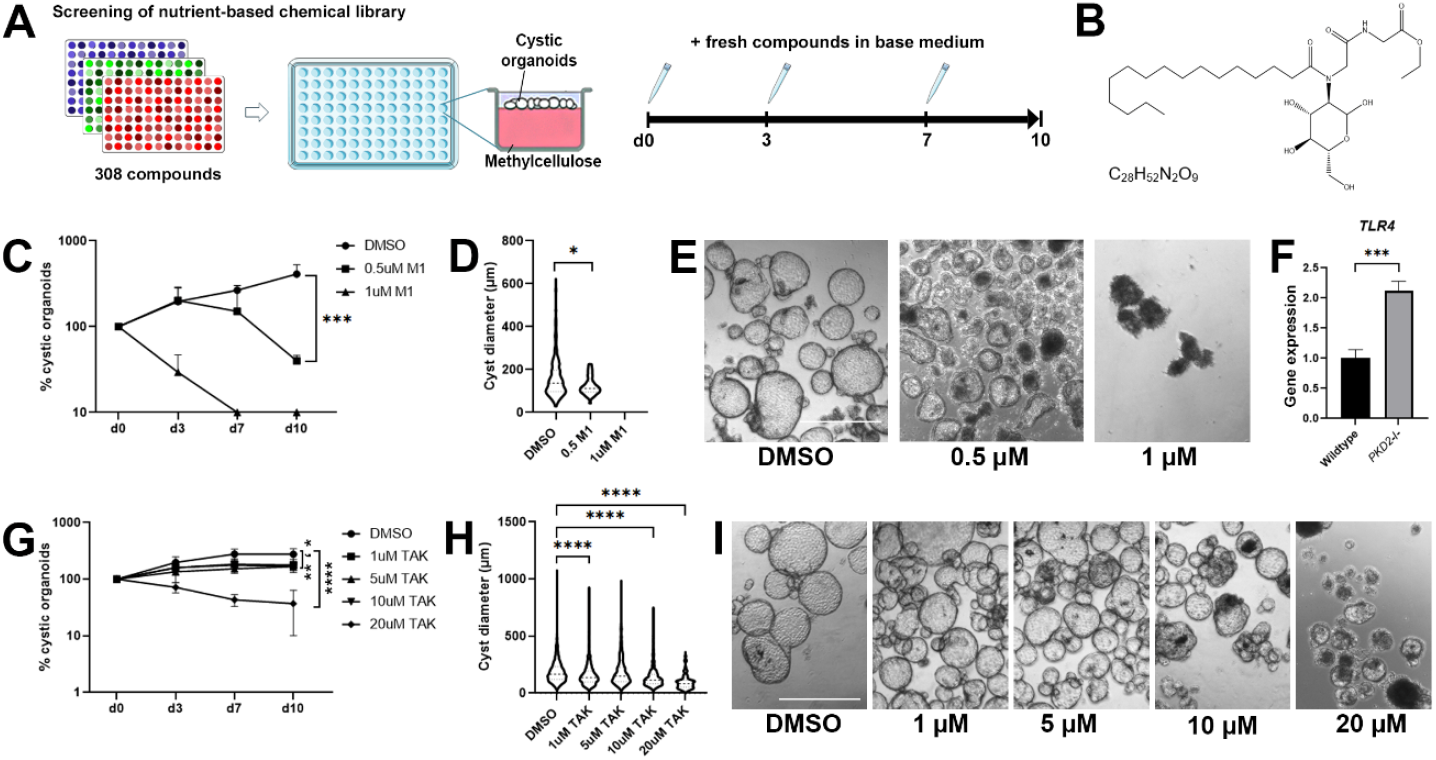
Library screening on methylcellulose-embedded cystic organoids. **A**. Schematic and timeline of the assay. **B**. Chemical structure of compound M1. **C**. Percent of cystic organoids after treatment with DMSO and different concentrations of M1 over 10 days. *** p≤ 0.001 (two-way ANOVA with Tukey’s multiple comparisons test). **D**. Cyst diameters after treatment with DMSO and M1 on day 10. * p≤ 0.05 (one-way ANOVA with Dunnett’s multiple comparisons test). **E**. Representative brightfield images of DMSO- and M1-treated UB *PKD2*^*-/-*^ organoids on day 10. Scale bar, 500 µm. **F**. Expression of *TLR4* in wildtype and *PKD2*^*-/-*^ UB organoids. *** p≤ 0.001 (Unpaired *t*-test) **G**. Percent of cystic organoids after treatment with DMSO and different concentrations of TAK-242 over 10 days. * p≤ 0.05 (1 µM); ** p≤ 0.01 (5 and 10 µM); **** p≤ 0.0001 (20 µM; two-way ANOVA with Tukey’s multiple comparisons test). **H**. Cyst diameters after treatment with DMSO and TAK-242 on day 10. **** p≤ 0.0001 (one-way ANOVA with Dunnett’s multiple comparisons test). **I**. Representative brightfield images of DMSO- and TAK-242-treated UB *PKD2*^*-/-*^ organoids on day 10. Scale bar, 500 µm.

## DISCUSSION

The aim of this study was to create a human organoid model of PKD that allows for drug screening. We found that switching from nephron organoids to UB organoids and growing these in suspension was key for generating large cyst cultures. Our UB organoid protocol relies on the transdifferentiation potential of the distal nephron cells within nephron organoids into UB epithelia^7^. Suspension culture throughout our protocol results in the formation of small UB organoids that have a more compact shape and display less obvious branching compared to protocols that grow UB organoids embedded in matrix and/or on transwell membranes^7,10,11,19^. Our method can produce a large yield of organoids (up to 2000 organoids per assay) that retain a UB-specific gene signature^27^. We recreated the *PKD2* mutation previously shown to induce cysts in nephron organoids^8^. In our UB organoids, this leads to a rapid formation of budding cysts with close to 100% efficiency. Unlike other systems wherein the removal of adherent cues or treatment with forskolin are needed to promote cystogenesis, the cysts in our method form spontaneously and express markers of ADPKD-associated YAP, JAK-STAT and NF-κB pathway activation, as well as early markers of fibrosis.

We validated our platform against known cyst-inhibitory compounds, including JQ1, QNZ and tolvaptan and found effective cyst reduction with QNZ^6^, although QNZ toxicity was observed at a lower concentration in our organoids (UB and nephron) than in the previous study, possible due to differences in the protocols. We found that the BET inhibitor JQ1 significantly reduces cysts in our UB organoids, which validates its therapeutic effects in *Pkd1* mutant mice^30^. BET inhibitors disrupt the interactions between bromodomain proteins and chromatin and are thus used to epigenetically modulate gene expression associated with aberrant cell growth and other pathogenic processes^39^. Brd4 protein is found upregulated in *Pkd1*^-/-^ mice and JQ1 treatment caused a decrease in c-Myc levels, phosphorylation of Rb and proliferation of the cystic epithelia. A similar mechanism could be responsible for the cyst reduction observed in our organoids, as some of the cystic epithelial cells can be stained for KI67, a marker for cell division (data not shown). BET proteins also regulate NF-κB associated inflammation and fibrosis^40–42^. NF-κB is constitutively activated in ADPKD and it has been suggested to be a key contributor to the inflammatory environment in cystic kidneys that promotes disease progression^43^. Thus, targeting NF-κB signaling with QNZ or JQ1 may be a promising strategy for PKD therapy.

Tolvaptan acts by lowering the elevated levels of intracellular cAMP, a hallmark of ADPKD cysts, via the inhibition of the vasopressin V2 receptor. Previous studies using hPSC-derived PKD organoids found that organoid cysts were unresponsive to tolvaptan^6,11,12^. Likewise, we did not observe cyst reduction with tolvaptan in our *PKD2*^*-/-*^ UB organoids. In contrast, primary cyst cultures derived from ADPKD patient kidneys respond to 10 nM tolvaptan^44^. Similarly, cyst-forming tubuloids derived from *PKD1* or *PKD2*-mutant adult stem cells respond to tolvaptan with cyst reduction, however at a higher concentration (15 µM) than the primary human cyst cultures^45^. Both these adult kidney-derived *in vitro* cultures require vasopressin stimulation for robust cyst formation, demonstrating the presence of functional vasopressin receptors. The study utilizing tubuloids also found that *AVPR2* expression was highly increased compared to iPSC-derived kidney organoids. From gene and protein expression analysis it is known that hPSC-derived kidney organoids are not fully differentiated and resemble human kidney tissue in the first trimester^21,46,47^. In line with this, we observed ∼3-times lower *AVPR2* expression in UB organoids than in human fetal and adult kidney samples, indicating that immature expression levels of vasopressin receptor likely account for the ineffectiveness of tolvaptan in the organoid systems. Similarly, our organoids are not exposed to tubular flow, which has been shown to improve organoid maturation and gene expression^48^. Hiratsuka *et al*^13^ recently showed cyst reduction with the FOS/AP1 inhibitor T-5224 in *PKHD1*^*-/-*^ nephron organoids under flow. In contrast, our *PKD2*^*-/-*^ UB organoids grown in shaking suspension culture responded to T-5224 treatment by cyst expansion, highlighting the differences between the two organoid models.

With the need for better therapies than tolvaptan, we tested the applicability of our UB organoid platform for drug discovery with a proof-of-principle library screen and identified compound M1 to be effective in reducing cysts. Compound M1 is composed of one chain of palmitate, one glucosamine, and two glycines. The structure closely resembles the natural bacterial molecule N-palmitoyl-D-glucosamine (PGA). PGA has been shown to antagonize TLR4 signaling *in vivo*, preventing lipopolysaccharide (LPS)-induced inflammation and neuropathic pain and reducing colon inflammation in mice^34,35^. The lipid A component of LPS, which is composed of two glucosamines and six fatty acid chains, is a strong inducer of the innate immune response via TLR4, whereas PGA and other under-acetylated variants of lipid A act as TLR4 antagonists^49^. Based on this, M1 may act as a TLR4 antagonist and given that the TLR4 signaling pathway operates via NF-κB, its inhibitory effects on cystogenesis may be via a similar mechanism to QNZ. The TLR4 pathway is relatively underexplored in PKD. In *Pkd1*^*RC/RC*^ mice, *Tlr4* is upregulated in the kidney as shown by RNA-seq, but this does not distinguish between immune cells and the renal parenchyma^50^. A single cell RNA-seq analysis of human ADPKD kidneys found that *TLR4* is highly expressed in leukocytes^27^ while other work has linked its expression to disease progression^51^. Increased *TLR4* expression in the renal epithelia has been reported in ischemic and nephrotoxic acute kidney injury^52,53^. Less is known about TLR4 in cystic epithelia, but TLR4 co-receptor CD14 was shown localized to cystic renal tubular epithelia and correlating to disease progression in *cpk* mice^54^, suggesting a role of TLR4 signaling in PKD. Kidney organoids lack an immune system so they will be a useful tool to address this knowledge gap. We show that *TLR4* expression is upregulated in cystic *PKD2*^*-/-*^ UB organoids and that TLR4 inhibition with TAK-242 reduces cyst numbers and size. A similar response was reported in *Pkd1*^*RC/RC*^ and *Pkd2*-KO mice, where miR-214 was upregulated in the interstitium of the cyst microenvironment and in turn inhibited *Tlr4* expression, thus protecting from inflammation and cyst growth^55^. These observations raise the possibility that TLR4 activation in UB/collecting duct epithelia or interstitial cells may play a key non-immune-related role in the early stages of cyst initiation in ADPKD^56^, and that targeting of TLR4 could be an effective strategy for cyst reduction. In summary, we present a protocol for generating UB organoids in suspension culture that is highly efficient for cyst production and amenable to small molecule screening.

## ACKNOWLEDGEMENTS

This work was supported by the China-Maurice Wilkins Centre project grant to AJD’s lab, and in part by JSPS (22H00350) to MU’s lab.

## AUTHOR CONTRIBUTIONS

VS and AJD conceived and designed the research; RW, TV and VS performed the experiments, analyzed the data and interpreted the results; RW and VS prepared the figures; MU and MT provided the drug library; VS wrote the manuscript; All authors edited and revised the manuscript; All authors approved the final version of manuscript.

## SUPPLEMENTARY MATERIAL

**Supplementary Figure S1**. Scalability of the UB organoid assay

**Supplementary Figure S2**. Efficiency of cyst formation in nephron and UB organoids

**Supplementary Figure S3**. Toxicity on nephron organoids

**Supplementary Figure S4**. FOS/AP1 inhibitor T-5224 increases cyst size

**Supplementary Table S1**. List of qPCR primers

**Figure S1.**
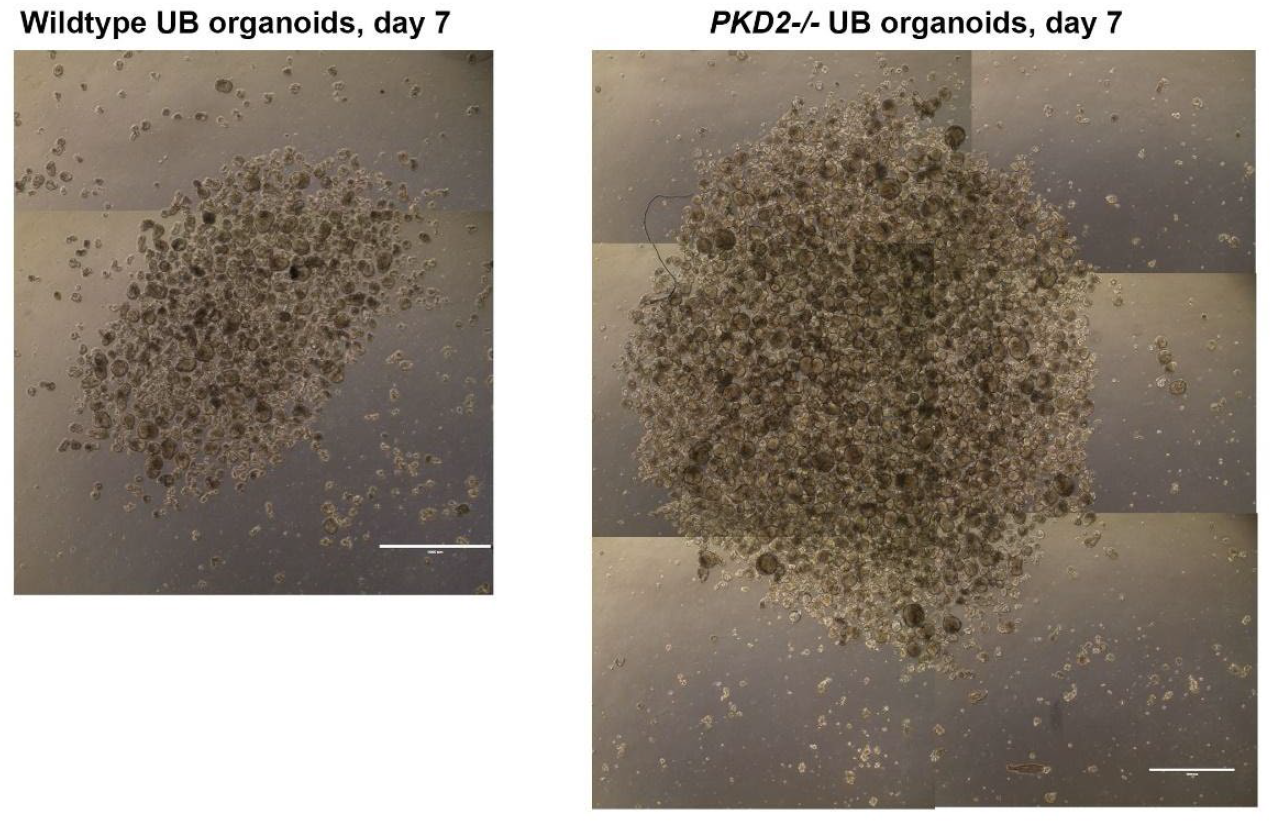
Scalability of the UB organoid assay. Representative brightfield images of single 6-wells of wildtype and *PKD2*^*-/-*^ UB organoid assays on day 7. Scale bars, 1000 µm.

**Figure S2.**
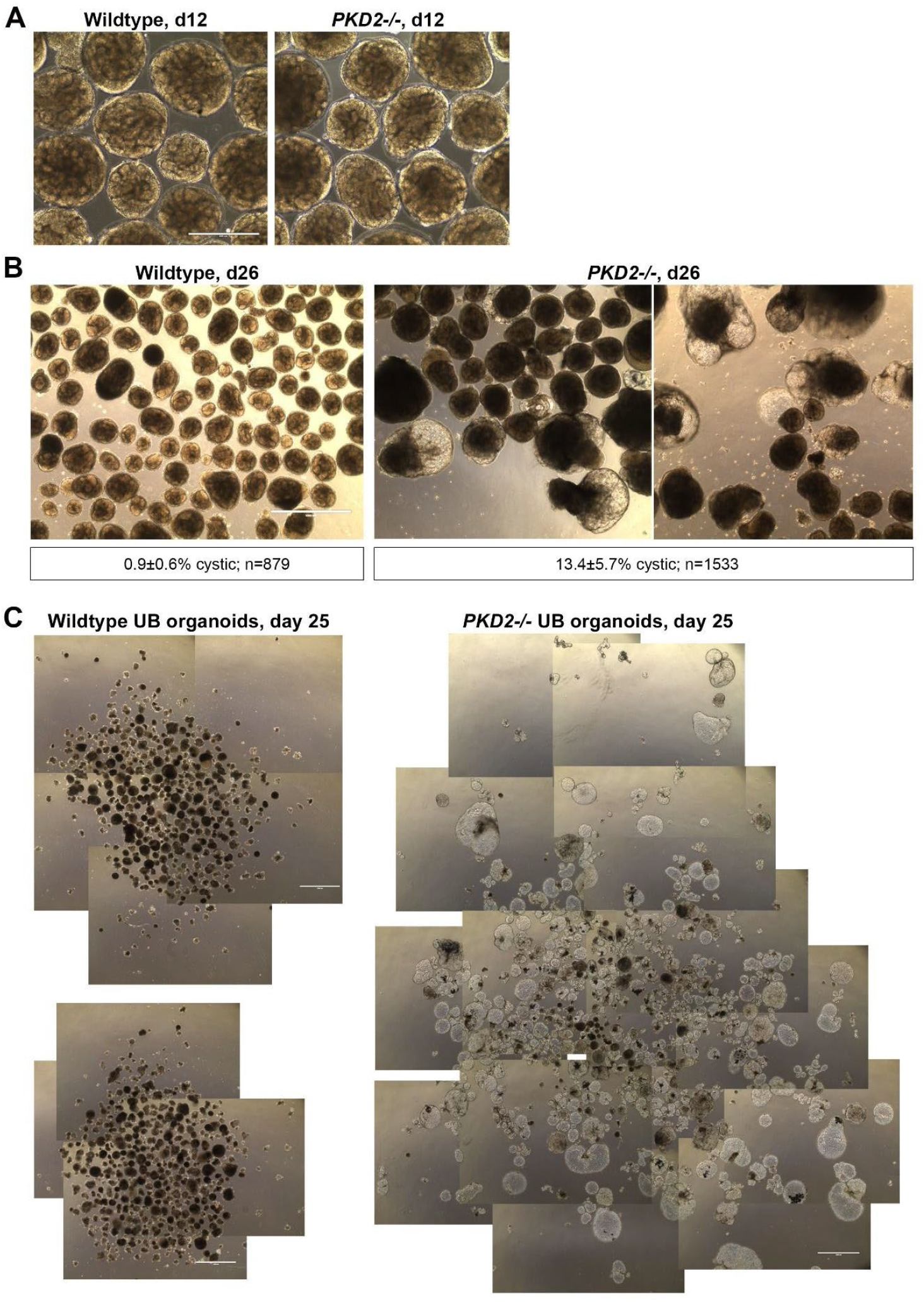
Efficiency of cyst formation in nephron and UB organoids. **A**. Wildtype and *PKD2*^*-/-*^ nephron organoids develop indistinguishably. **B**. Cyst formation in wildtype and *PKD2*^*-/-*^ nephron organoids. **C**. Wildtype (two separate 6-wells) and *PKD2*^*-/-*^ UB organoid assays on day 25 showing expansion of the cystic organoid assay. Scale bars, 400 µm (A); 1000 µm (B, C).

**Figure S3.**
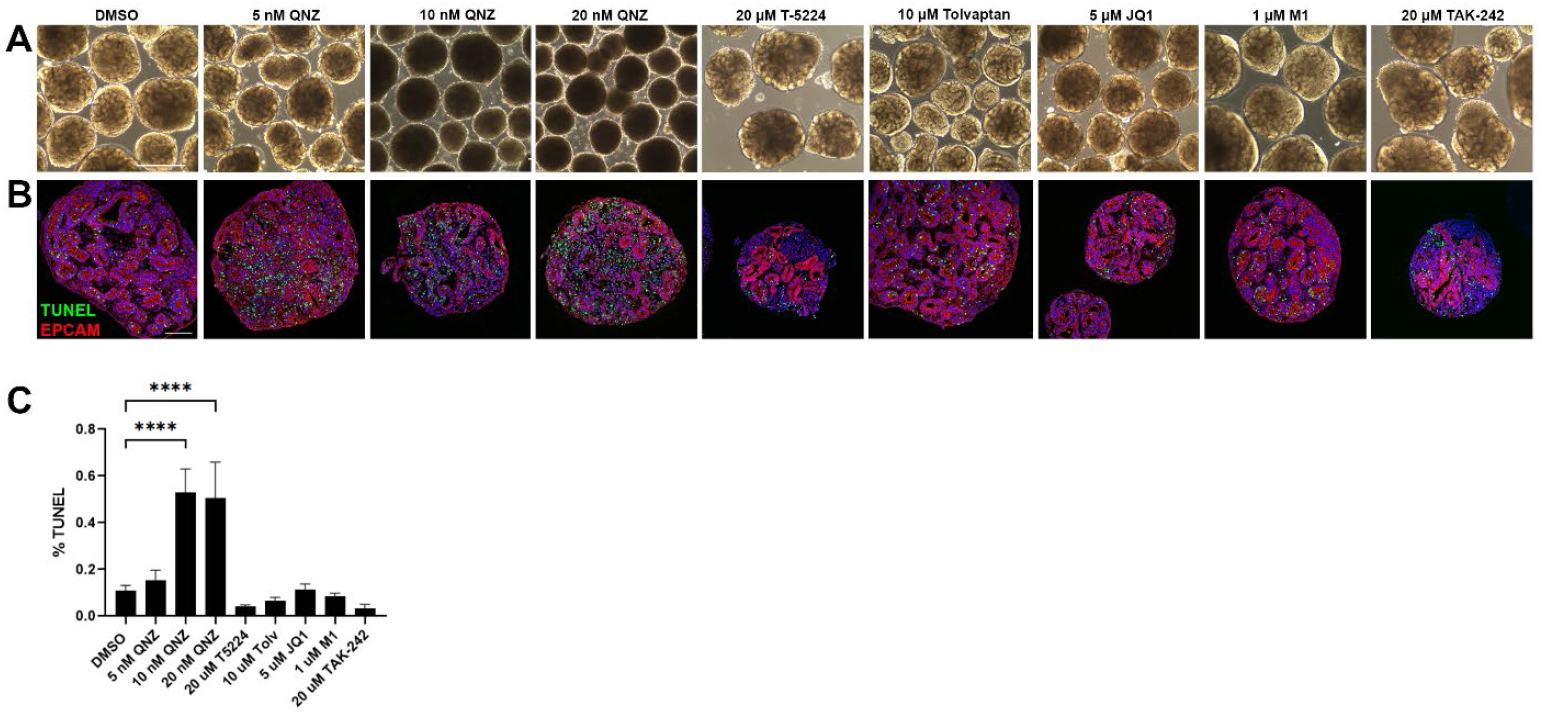
Toxicity on nephron organoids. **A**. Brightfield images of nephron organoids after treatment with DMSO, QNZ, T-5224, tolvaptan, JQ1, M1, and TAK-242 for 72 hrs. **B**. Paraffin sections of nephron organoids labeled for cell death (TUNEL, green) and tubules (EPCAM, red). Nuclear counterstain, Hoechst. **C**. Quantification of TUNEL-positive cells. n≥10 sections per condition. **** p≤ 0.0001 (one-way ANOVA with Dunnett’s multiple comparisons test). Scale bars, 400 µm (A); 100 µm (B).

**Figure S4.**
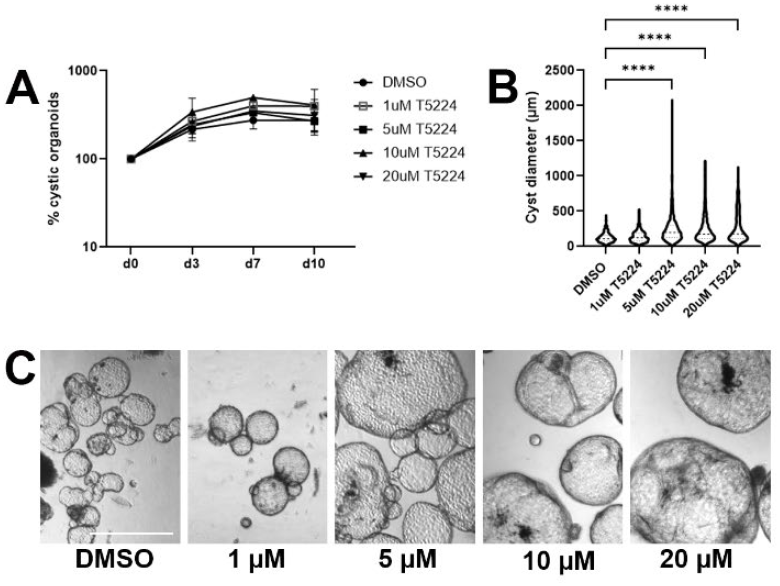
FOS/AP1 inhibitor T-5224 increases cyst size. **A**. Percent of cystic organoids after treatment with DMSO and different concentrations of T-5224 over 10 days. **B**. Cyst diameters after treatment with DMSO and T-5224 on day 10. **** p≤ 0.0001 (one-way ANOVA with Dunnett’s multiple comparisons test). **C**. Representative brightfield images of DMSO- and T-5224-treated UB *PKD2*^*-/-*^ organoids on day 10. Scale bar, 500 µm.

**Table S1.**
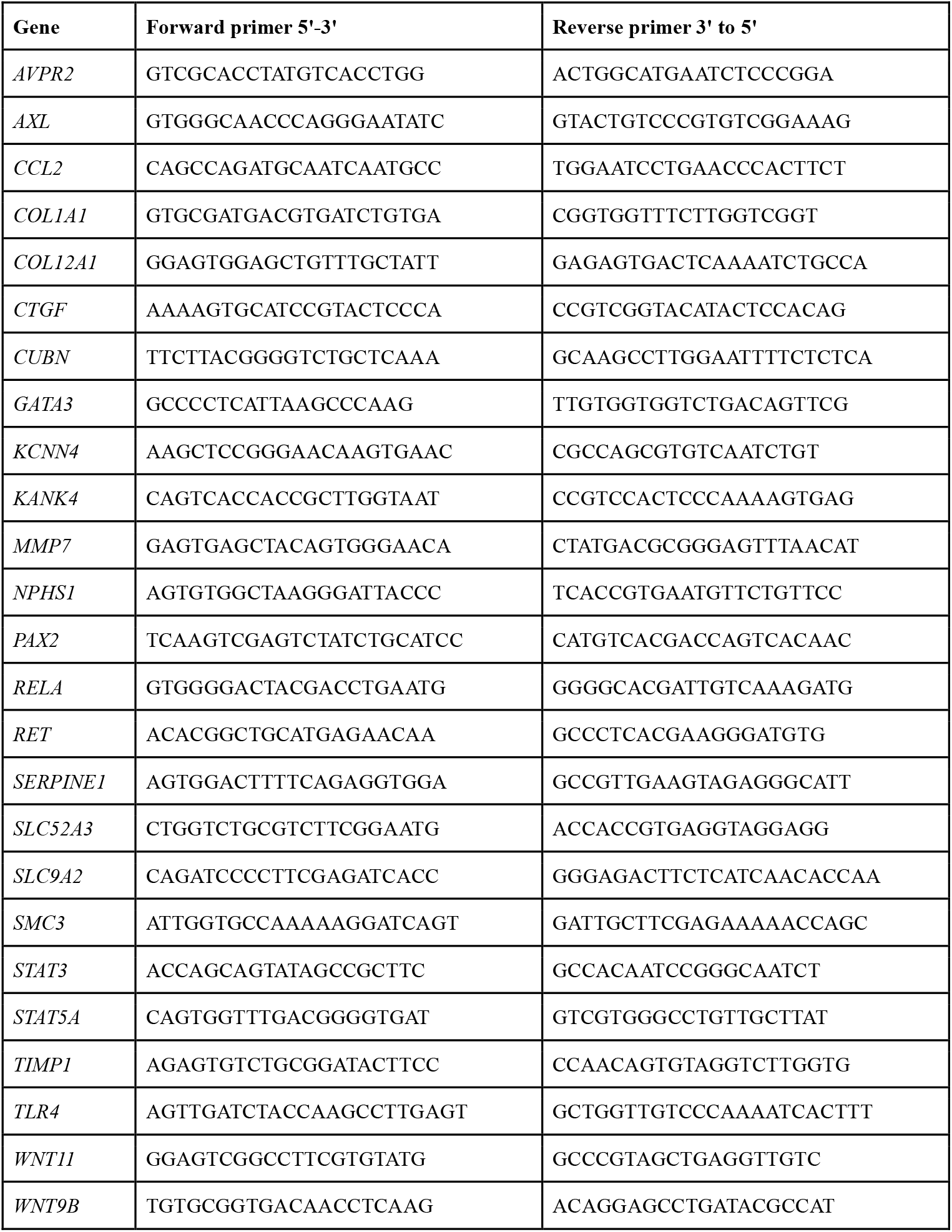
List of qPCR primers.

## REFERENCES

1. Bergmann C, Guay-Woodford LM, Harris PC, Horie S, Peters DJM, Torres VE. Polycystic kidney disease. Nat Rev Dis Primers. 2018;4(1):50. doi:10.1038/s41572-018-0047-y

2. Raina R, Houry A, Rath P, et al. Clinical Utility and Tolerability of Tolvaptan in the Treatment of Autosomal Dominant Polycystic Kidney Disease (ADPKD). Drug, Healthc Patient Saf. 2022;14:147–159. doi:10.2147/dhps.s338050

3. Leonhard WN, Happe H, Peters DJM. Variable Cyst Development in Autosomal Dominant Polycystic Kidney Disease: The Biologic Context. J Am Soc Nephrol. 2016;27(12):3530–3538. doi:10.1681/asn.2016040425

4. Grantham JJ, Mulamalla S, Swenson-Fields KI. Why kidneys fail in autosomal dominant polycystic kidney disease. Nat Rev Nephrol. 2011;7(10):556–566. doi:10.1038/nrneph.2011.109

5. Barnawi RA, Attar RZ, Alfaer SS, Safdar OY. Is the light at the end of the tunnel nigh? A review of ADPKD focusing on the burden of disease and tolvaptan as a new treatment. Int J Nephrol Renov Dis. 2018;11:53–67. doi:10.2147/ijnrd.s136359

6. Tran T, Song CJ, Nguyen T, et al. A scalable organoid model of human autosomal dominant polycystic kidney disease for disease mechanism and drug discovery. Cell Stem Cell. 2022;29(7):1083-1101.e7. doi:10.1016/j.stem.2022.06.005

7. Howden SE, Wilson SB, Groenewegen E, et al. Plasticity of distal nephron epithelia from human kidney organoids enables the induction of ureteric tip and stalk. Cell Stem Cell. Published online 2020. doi:10.1016/j.stem.2020.12.001

8. Freedman BS, Brooks CR, Lam AQ, et al. Modelling kidney disease with CRISPR-mutant kidney organoids derived from human pluripotent epiblast spheroids. Nature Communications. 2015;6(1):ncomms9715. doi:10.1038/ncomms9715

9. Low JH, Li P, Chew EGY, et al. Generation of Human PSC-Derived Kidney Organoids with Patterned Nephron Segments and a De Novo Vascular Network. Cell Stem Cell. Published online 2019. doi:10.1016/j.stem.2019.06.009

10. Liu M, Zhang C, Gong X, et al. Kidney organoid models reveal cilium-autophagy metabolic axis as a therapeutic target for PKD both in vitro and in vivo. Cell Stem Cell. 2024;31(1):52-70.e8. doi:10.1016/j.stem.2023.12.003

11. Shimizu T, Mae SI, Araoka T, et al. A novel ADPKD model using kidney organoids derived from disease-specific human iPSCs. Biochemical and Biophysical Research Communications. 2020;(4). doi:10.1016/j.bbrc.2020.06.141

12. Huang B, Zeng Z, Kim S, et al. Long-term expandable mouse and human-induced nephron progenitor cells enable kidney organoid maturation and modeling of plasticity and disease. Cell Stem Cell. 2024;31(6):921-939.e17. doi:10.1016/j.stem.2024.04.002

13. Hiratsuka K, Miyoshi T, Kroll KT, et al. Organoid-on-a-chip model of human ARPKD reveals mechanosensing pathomechanisms for drug discovery. Sci Adv. 2022;8(38):eabq0866. doi:10.1126/sciadv.abq0866

14. Vishy CE, Thomas C, Vincent T, Crawford DK, Goddeeris MM, Freedman BS. Genetics of cystogenesis in base-edited human organoids reveal therapeutic strategies for polycystic kidney disease. Cell Stem Cell. 2024;31(4):537-553.e5. doi:10.1016/j.stem.2024.03.005

15. Taguchi A, Nishinakamura R. Higher-Order Kidney Organogenesis from Pluripotent Stem Cells. Cell Stem Cell. 2017;21(6):730-746.e6. doi:10.1016/j.stem.2017.10.011

16. Zeng Z, Huang B, Parvez RK, et al. Generation of patterned kidney organoids that recapitulate the adult kidney collecting duct system from expandable ureteric bud progenitors. Nat Commun. 2021;12(1):3641. doi:10.1038/s41467-021-23911-5

17. Shi M, McCracken KW, Patel AB, et al. Human ureteric bud organoids recapitulate branching morphogenesis and differentiate into functional collecting duct cell types. Nat Biotechnol. Published online 2022:1-10. doi:10.1038/s41587-022-01429-5

18. Mae SI, Ryosaka M, Sakamoto S, et al. Expansion of Human iPSC-Derived Ureteric Bud Organoids with Repeated Branching Potential. Cell Reports. 2020;32(4):107963. doi:10.1016/j.celrep.2020.107963

19. Kuraoka S, Tanigawa S, Taguchi A, et al. PKD1-Dependent Renal Cystogenesis in Human Induced Pluripotent Stem Cell-Derived Ureteric Bud/Collecting Duct Organoids. J Am Soc Nephrol Jasn. Published online 2020:ASN.2020030378. doi:10.1681/asn.2020030378

20. Mae SI, Hattanda F, Morita H, et al. Human iPSC-derived renal collecting duct organoid model cystogenesis in ADPKD. Cell Rep. 2023;42(12):113431. doi:10.1016/j.celrep.2023.113431

21. Przepiorski A, Sander V, Tran T, et al. A Simple Bioreactor-Based Method to Generate Kidney Organoids from Pluripotent Stem Cells. Stem Cell Reports. 2018;(2). doi:10.1016/j.stemcr.2018.06.018

22. Sander V, Przepiorski A, Crunk AE, Hukriede NA, Holm TM, Davidson AJ. Protocol for Large-Scale Production of Kidney Organoids from Human Pluripotent Stem Cells. Star Protoc. Published online 2020:100150. doi:10.1016/j.xpro.2020.100150

23. Oh JK, Przepiorski A, Chang HH, et al. Derivation of induced pluripotent stem cell lines from New Zealand donors. J Roy Soc New Zeal. Published online 2020:1-14. doi:10.1080/03036758.2020.1830808

24. Harris PWR, Siow A, Yang SH, et al. Synthesis, Antibacterial Activity, and Nephrotoxicity of Polymyxin B Analogues Modified at Leu-7, d-Phe-6, and the N-Terminus Enabled by S-Lipidation. Acs Infect Dis. Published online 2022. doi:10.1021/acsinfecdis.1c00347

25. Yuri S, Nishikawa M, Yanagawa N, Jo OD, Yanagawa N. In Vitro Propagation and Branching Morphogenesis from Single Ureteric Bud Cells. Stem Cell Rep. 2017;8(2):401–416. doi:10.1016/j.stemcr.2016.12.011

26. Cruz NM, Song X, Czerniecki SM, et al. Organoid cystogenesis reveals a critical role of microenvironment in human polycystic kidney disease. Nature Materials. 2017;16(11):1112–1119. doi:10.1038/nmat4994

27. Muto Y, Dixon EE, Yoshimura Y, et al. Defining cellular complexity in human autosomal dominant polycystic kidney disease by multimodal single cell analysis. Nat Commun. 2022;13(1):6497. doi:10.1038/s41467-022-34255-z

28. Formica C, Peters DJM. Molecular pathways involved in injury-repair and ADPKD progression. Cell Signal. 2020;72:109648. doi:10.1016/j.cellsig.2020.109648

29. Uchihashi S, Fukumoto H, Onoda M, Hayakawa H, Ikushiro S ichi, Sakaki T. Metabolism of the c-Fos/Activator Protein-1 Inhibitor T-5224 by Multiple Human UDP-Glucuronosyltransferase Isoforms. Drug Metab Dispos. 2011;39(5):803–813. doi:10.1124/dmd.110.037952

30. Zhou X, Fan LX, Peters DJM, Trudel M, Bradner JE, Li X. Therapeutic targeting of BET bromodomain protein, Brd4, delays cyst growth in ADPKD. Hum Mol Genet. 2015;24(14):3982–3993. doi:10.1093/hmg/ddv136

31. Furuta T, Mizukami Y, Asano L, et al. Nutrient-Based Chemical Library as a Source of Energy Metabolism Modulators. Acs Chem Biol. 2019;14(9):1860–1865. doi:10.1021/acschembio.9b00444

32. Vale N, Ferreira A, Matos J, Fresco P, Gouveia MJ. Amino Acids in the Development of Prodrugs. Molecules. 2018;23(9):2318. doi:10.3390/molecules23092318

33. Jaracz S, Chen J, Kuznetsova LV, Ojima I. Recent advances in tumor-targeting anticancer drug conjugates. Bioorg Med Chem. 2005;13(17):5043–5054. doi:10.1016/j.bmc.2005.04.084

34. Iannotta M, Belardo C, Trotta MC, et al. N-palmitoyl-D-glucosamine, A Natural Monosaccharide-Based Glycolipid, Inhibits TLR4 and Prevents LPS-Induced Inflammation and Neuropathic Pain in Mice. Int J Mol Sci. 2021;22(3):1491. doi:10.3390/ijms22031491

35. Palenca I, Seguella L, Re AD, et al. N-Palmitoyl-D-Glucosamine Inhibits TLR-4/NLRP3 and Improves DNBS-Induced Colon Inflammation through a PPAR-α-Dependent Mechanism. Biomolecules. 2022;12(8):1163. doi:10.3390/biom12081163

36. Ii M, Matsunaga N, Hazeki K, et al. A Novel Cyclohexene Derivative, Ethyl (6R)-6-[N-(2-Chloro-4-fluorophenyl)sulfamoyl]cyclohex-1-ene-1-carboxylate (TAK-242), Selectively Inhibits Toll-Like Receptor 4-Mediated Cytokine Production through Suppression of Intracellular Signaling. Mol Pharmacol. 2006;69(4):1288–1295. doi:10.1124/mol.105.019695

37. Fenhammar J, Rundgren M, Forestier J, Kalman S, Eriksson S, Frithiof R. Toll-Like Receptor 4 Inhibitor TAK-242 Attenuates Acute Kidney Injury in Endotoxemic Sheep. Anesthesiology. 2011;114(5):1130–1137. doi:10.1097/aln.0b013e31820b8b44

38. Rice TW, Wheeler AP, Bernard GR, et al. A randomized, double-blind, placebo-controlled trial of TAK-242 for the treatment of severe sepsis & Crit Care Med. 2010;38(8):1685–1694. doi:10.1097/ccm.0b013e3181e7c5c9

39. To KKW, Xing E, Larue RC, Li PK. BET Bromodomain Inhibitors: Novel Design Strategies and Therapeutic Applications. Molecules. 2023;28(7):3043. doi:10.3390/molecules28073043

40. Xiong C, Masucci MV, Zhou X, et al. Pharmacological targeting of BET proteins inhibits renal fibroblast activation and alleviates renal fibrosis. Oncotarget. 2016;7(43):69291–69308. doi:10.18632/oncotarget.12498

41. Morgado-Pascual JL, Rayego-Mateos S, Tejedor L, Suarez-Alvarez B, Ruiz-Ortega M. Bromodomain and Extraterminal Proteins as Novel Epigenetic Targets for Renal Diseases. Front Pharmacol. 2019;10:1315. doi:10.3389/fphar.2019.01315

42. Hajmirza A, Emadali A, Gauthier A, Casasnovas O, Gressin R, Callanan MB. BET Family Protein BRD4: An Emerging Actor in NFκB Signaling in Inflammation and Cancer. Biomedicines. 2018;6(1):16. doi:10.3390/biomedicines6010016

43. Ta MH, Schwensen KG, Liuwantara D, Huso DL, Watnick T, Rangan GK. Constitutive renal Rel/nuclear factor-κB expression in Lewis polycystic kidney disease rats. World J Nephrol. 2016;5(4):339. doi:10.5527/wjn.v5.i4.339

44. Reif GA, Yamaguchi T, Nivens E, Fujiki H, Pinto CS, Wallace DP. Tolvaptan inhibits ERK-dependent cell proliferation, Cl− secretion and in vitro cyst growth of human ADPKD cells stimulated by vasopressin. Am J Physiol-renal. 2011;301(5):F1005–F1013. doi:10.1152/ajprenal.00243.2011

45. Xu Y, Kuppe C, Perales-Patón J, et al. Adult human kidney organoids originate from CD24+ cells and represent an advanced model for adult polycystic kidney disease. Nat Genet. 2022;54(11):1690–1701. doi:10.1038/s41588-022-01202-z

46. Takasato M, Er PX, Chiu HS, et al. Kidney organoids from human iPS cells contain multiple lineages and model human nephrogenesis. Nature. 2015;526(7574):nature15695. doi:10.1038/nature15695

47. Combes AN, Zappia L, Er PX, Oshlack A, Little MH. Single-cell analysis reveals congruence between kidney organoids and human fetal kidney. Genome Med. 2019;11(1):3. doi:10.1186/s13073-019-0615-0

48. Homan KA, Gupta N, Kroll KT, et al. Flow-enhanced vascularization and maturation of kidney organoids in vitro. Nat Methods. 2019;(3):1–8. doi:10.1038/s41592-019-0325-y

49. Heine H, Zamyatina A. Therapeutic Targeting of TLR4 for Inflammation, Infection, and Cancer: A Perspective for Disaccharide Lipid A Mimetics. Pharmaceuticals. 2022;16(1):23. doi:10.3390/ph16010023

50. Swenson-Fields KI, Ward CJ, Lopez ME, et al. Caspase-1 and the inflammasome promote polycystic kidney disease progression. Front Mol Biosci. 2022;9:971219. doi:10.3389/fmolb.2022.971219

51. Kocyigit I, Sener EF, Taheri S, et al. Toll-Like Receptors in the Progression of Autosomal Dominant Polycystic Kidney Disease. Ther Apher Dial. 2016;20(6):615–622. doi:10.1111/1744-9987.12458

52. Wu H, Chen G, Wyburn KR, et al. TLR4 activation mediates kidney ischemia/reperfusion injury. J Clin Investig. 2007;117(10):2847–2859. doi:10.1172/jci31008

53. Zhang B, Ramesh G, Uematsu S, Akira S, Reeves WB. TLR4 Signaling Mediates Inflammation and Tissue Injury in Nephrotoxicity. J Am Soc Nephrol. 2008;19(5):923–932. doi:10.1681/asn.2007090982

54. Zhou J, Ouyang X, Cui X, et al. Renal CD14 expression correlates with the progression of cystic kidney disease. Kidney Int. 2010;78(6):550–560. doi:10.1038/ki.2010.175

55. Lakhia R, Yheskel M, Flaten A, et al. Interstitial microRNA miR-214 attenuates inflammation and polycystic kidney disease progression. JCI Insight. 2020;5(7). doi:10.1172/jci.insight.133785

56. Anders HJ, Banas B, Schlöndorff D. Signaling Danger & Toll-Like Receptors and their Potential Roles in Kidney Disease. J Am Soc Nephrol. 2004;15(4):854–867. doi:10.1097/01.asn.0000121781.89599.16

